# High-order interactions dominate the functional landscape of microbial consortia

**DOI:** 10.1101/333534

**Authors:** Alicia Sanchez-Gorostiaga, Djordje Bajić, Melisa L. Osborne, Juan F. Poyatos, Alvaro Sanchez

**Author notes:** These authors contributed equally (Alicia Sanchez-Gorostiaga, Djordje Bajić). (JFP), (AS).

## Abstract

Understanding the link between community composition and function is a major challenge in microbial ecology, with implications for the management of natural microbiomes and the design of synthetic consortia. For this purpose, it is critical to understand the extent to which community functions and properties can be predicted from species traits and what role is played by complex interactions. Inspired by the study of complex genetic interactions and fitness landscapes, here we have examined how the amylolytic function of combinatorial assemblages of seven starch-degrading soil bacteria depends on the functional contributions from each species and their interactions. Filtering our experimental results through the theory of enzyme kinetics, we show that high-order functional interactions dominate the amylolytic rate of our consortia, even though this function is biochemically simple, redundantly distributed in the community, and additive in the absence of inter-species interactions. As the community grows in size, the contribution of high-order functional interactions grows too, making the community function increasingly unpredictable. We can explain the prevalence of high order effects and their sign from the redundancy of ecological interactions in the network, in particular from redundant facilitation towards a high-performing community member. Our results suggest that even simple functions can be dominated by complex interactions, posing challenges for the predictability and bottom-up engineering of ecosystem function in complex multi-species communities.

## Introduction

Microbial communities carry out critical biochemical functions throughout the biosphere: from nitrogen fixation and photosynthesis to the recycling of nutrients and the decomposition of inert organic matter [1,2]. In host associated communities, the metabolic activity of the microbiota can also profoundly affect the host’s health, and has been found to modulate life history traits such as flowering timing in plants [3,4] or the lifespan and reproductive behavior of animals [5,6]. In more applied settings, microbial consortia are being designed for the processing of undesirable materials into valuable products [7–11], to protect valuable crops from pathogens [12,13] or other stressors [14,15], or even to increase crop yields [16]. All of these effects that microbial communities have on the environment and their hosts are often referred to as “functions” of these communities. These functions depend very strongly on community composition: i.e. on which species are present and in what numbers. Thus, manipulating community composition to accomplish desirable functional outcomes has become a major goal in fields as diverse as medicine, environmental engineering, and biotechnology [15].

To accomplish this goal, it is imperative to develop a predictive understanding of the relationship between microbial community composition and function [17,18]. This is widely recognized as one of the main challenges in the field [7,17–21], but many fundamental questions remain. Importantly, it is still unclear to what extent one can predict the function of a large multi-species community from low-dimensional information, such as the functional contributions of single species and their pairwise interactions [22–26]. The answer to this question has important implications: If predicting community function in such a bottom-up manner were generally feasible, this would encourage synthetic approaches to designing complex communities in a rational manner, by mixing and matching components with known functional traits [7,18,26,27]. However, the contributions of a given species or pair of species to a community function may also depend on the presence or absence of other taxa, for instance through ecological interactions that modulate species abundance. This can easily lead to higher-than-pairwise functional interactions. If community functions were generally complex and enriched in nonlinear or higher order functional interactions, bottom-up prediction of community function would be significantly more challenging. In this scenario, top-down approaches such as community-level selection [28] might be a more promising avenue for the manipulation and design of complex microbiomes [4,14,28–32].

The crux of the problem is thus the contribution of high order interactions to community functions. In spite of an increasing appreciation of the role that high-order interactions may play in ecosystems [25,33–35], we still know little about their contribution to specific processes and community-level functions [6,36]. In order to tackle this question, one would need to disentangle the contributions of single species in isolation from the effects of pairwise and higher order interactions. This is a notoriously challenging problem, particularly for complex functions for which “first principles” mechanistic models are hard to produce [37]. A similar problem has been encountered many times before in other areas of biology, most notably in the study of fitness landscapes and genetic interactions (i.e. epistasis) among mutations [38–40]. Typically, a model-free statistical approach is typically followed in these studies in order to detect and quantify interactions when mechanistic models are not available. In this model-free approach, one compares the quantitative phenotype of a multiple mutant with the expectation from a null model that assumes that no interactions exist; typically null models assume that the phenotypic effects of each mutation add up [41–43]. The difference between the measured phenotype and the null model prediction is attributed to genetic interactions [41,42]. This approach is routinely applied to empirical fitness landscapes to detect pairwise epistasis [38,42] as well as to determine the impact of high-order gene interactions on cellular phenotypes [37,39,40]. In recent years, this approach has also been extended to detect interactions in other complex biological systems, e.g. between transcription factors in combinatorial gene regulation [44,45], or among multiple drugs in antibiotic or cancer drug cocktails [46,47].

Applied to microbial communities, this strategy would involve reconstituting all possible combinatorial sub-communities (i.e. every possible monoculture as well as every possible pairwise co-culture, three species co-culture, four member co-culture etc.) and measuring their function; then comparing this measurement with the prediction from a null model to identify interactions [6,36]. Clearly, constructing a null model that captures the absence of interactions is critical for unambiguously establishing the contribution of both pairwise and high order interactions to community-level functions. Yet, for complex functions that are also often inherently non-linear and depend in poorly understood ways on the molecular interplay between hosts and microbes, it is far from obvious what that null model should be. Without it, the relative role of high-order interactions and the predictability of community function cannot be unequivocally determined. Here, we have tackled this problem by focusing on a simple community-level function that can be quantitatively modeled on solid biochemical grounds. This allows us to formulate a null model that is based on the theory of enzymatic kinetics and validate it experimentally. Armed with that, we are able to unambiguously show not only that functional interactions exist, but that high-order interactions often dominate the functional landscape of our microbial consortia. As we will show, our combination of theory and experiments also allows us to understand the ecological basis for these high-order functional interactions.

## Results

### A simple additive function in a simple in vitro consortium

We seeked to study a small synthetic consortium that could be combinatorially reconstituted *in vitro*, and which performs a simple function that could be mechanistically and quantitatively understood from first principles when interactions among species are absent. Having a quantitative mechanistic model would allow us to directly detect interactions, as deviations from the predicted effect of independent species contributions. This is important because many complex biochemical functions can be non-additive even in the absence of interactions [44].

For this end, we constructed a synthetic consortium made up by seven amylolytic soil bacteria: *Bacillus subtilis, Bacillus megaterium, Bacillus mojavensis, Paenibacillus polymyxa, Bacillus thuringiensis, Bacillus licheniformis*, and *Bacillus cereus* (Fig. 1A). As a simple function that can be mechanistically understood from first principles, we chose the starch hydrolysis rate of the enzymes released by the consortium (Fig. 1B-C). This function was expressed on agar plates as well as in liquid culture (Fig. 1A-D) by all members of the consortia in isolation (Fig. 1D). The amylolytic community function is thus redundantly distributed, and it is also a simple function that requires just one kind of extracellular enzyme (an endoamylase) on its biochemical pathway.

**Figure 1.**
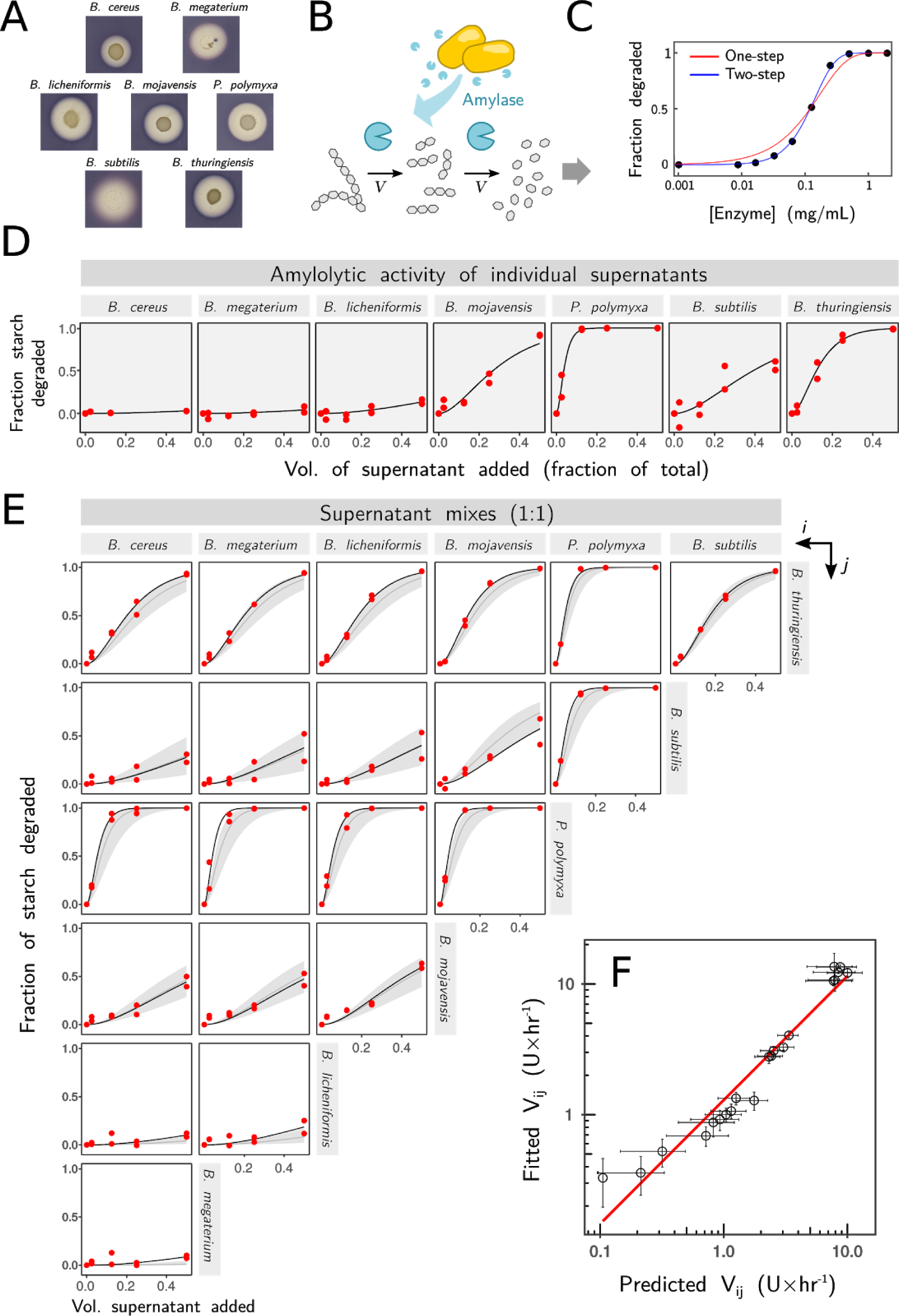
Starch degradation of simple consortia follows an additive null biochemical model. **A**.Extracellular degradation of starch is observed as a halo around colonies of different bacilli species growing on basic growth minimal media (1x bSAM). Starch is stained in blue using Lugol Iodine stain. **B.** Cartoon depicting the release of amylase (blue) and the extracellular degradation of starch (gray), which we model as a two-step process. **C.** Starch hydrolysis follows a sequential two-step reaction (black dots) that is well fitted by a two-step Michaelis-Menten Model. **D.** The kinetics of the amylolytic activity for the supernatants of individual species inoculated into 3mL of 1x bSAM and grown for 24hrs. We quantified the fraction of starch degraded by different dilutions of the filtered supernatants from each monoculture in reactions incubated at 30^°^C with 0.1% (w/v) starch for 180 min and quenched by a solution containing Lugol Iodine stain. Experimental data (red points) was fitted to Equation 2 (solid line, see Methods). **E.** Kinetics of the amylolytic activity for pairwise combinations (1:1) of the filtered supernatants of monocultures grown and assayed as in panel D. In addition to experimental data (red points) and fitted catalytic rate (*V*_*ij*_; solid black line), we also show the prediction from the individual models assuming catalytic rates of the individual enzymes are additive (gray line **±** 2SE). **F.** Comparison of fitted vs predicted values of *V*_*ij*_ for the same pairs shown in panel E (error bars represent **±** 2SE); red line represents perfect prediction. Note the log-log scale.

A kinetic *in vitro* measurement of starch hydrolysis by purified *B. subtilis* amylase (Fig. 1B-C) indicates that starch hydrolysis follows a sequential two-step reaction with two identical rates (*V*), which is well fitted by a Michaelis-Menten Model (Fig. 1C; See Methods). The same model also fits very well the kinetics of starch hydrolysis when we incubate soluble starch with the supernatants of every one of the species when grown independently in liquid culture (Fig. 1D). As shown in the Methods section, a kinetic model of enzymatic starch hydrolysis where the enzymes released by two species (*A* and *B*) act independently on the substrate predicts that the starch hydrolysis rate of (*V*_*AB*_) is the sum of the catalytic rates of each enzyme in isolation (*V*_*AB*_=*V*_*A*_+*V*_*B*_). We tested this prediction by growing all seven species in monocultures for 24 hours, and then mixing their supernatants in every possible pairwise combination (Fig. 1E). A series of different volumes of these supernatant admixtures were incubated with a 0.1% w/v starch solution for 2 hours, and the fraction of starch degraded in that time was then measured. A fit to the model in Eq. 2 (Methods) allowed us to obtain the catalytic rate of the enzymatic cocktails, which can be compared to the prediction from the model of independently acting enzymes (Fig. 1E). This naturally additive, interaction-free mechanistic model explains 97% of the variance observed when mixing every possible pair of enzymes from different species together (Fig. 1F).

In summary, the theory of chemical kinetics states that the rate of hydrolysis when two different enzymes are mixed together should be the sum of the hydrolysis rate of each enzyme, if those enzymes act independently on the substrate. This prediction is validated in our system. Therefore, when the species in our consortia are grown in separate culture tubes and thus do not interact with one another in any way, their contributions to the rate of starch degradation add up. We have thus selected a simple function that is additive in the absence of interactions, and which can be expressed redundantly by every member of our host-free consortium. This function can also be explained mechanistically in terms of simple biochemistry. Armed with this null mechanistic model, we are now set up to investigate the role of pairwise and higher order functional interactions in this microbial ecosystem.

### Pairwise functional interactions are ubiquitous in our microbial consortium

To investigate the contribution of pairwise and higher interactions to the function of our consortia we made use of tools that were originally developed for the study of genetic interactions. By analogy with Fitness Landscapes, we define the Functional Landscape as a map connecting every possible combinatorially assembled consortia with its function (which we generically denote as *F* and which, in the case at hand, it represents the amylolytic rate of the enzymes collectively secreted by the consortia). In the absence of interactions, and as predicted by the null model described above, the Functional Landscape is expected to be smooth, and the function would grow monotonically with the number of members of the consortia (Fig. 2A). To test this prediction, we assembled consortia formed by pairs, trios, as well as four, five, six and seven member species and measured the amylolytic rates of the enzyme cocktails they released after 24 hours of co-culture (Methods). The results (Fig. 2B) show a marked deviation from the null additive model, indicating the presence of substantial functional interactions.

**Figure 2.**
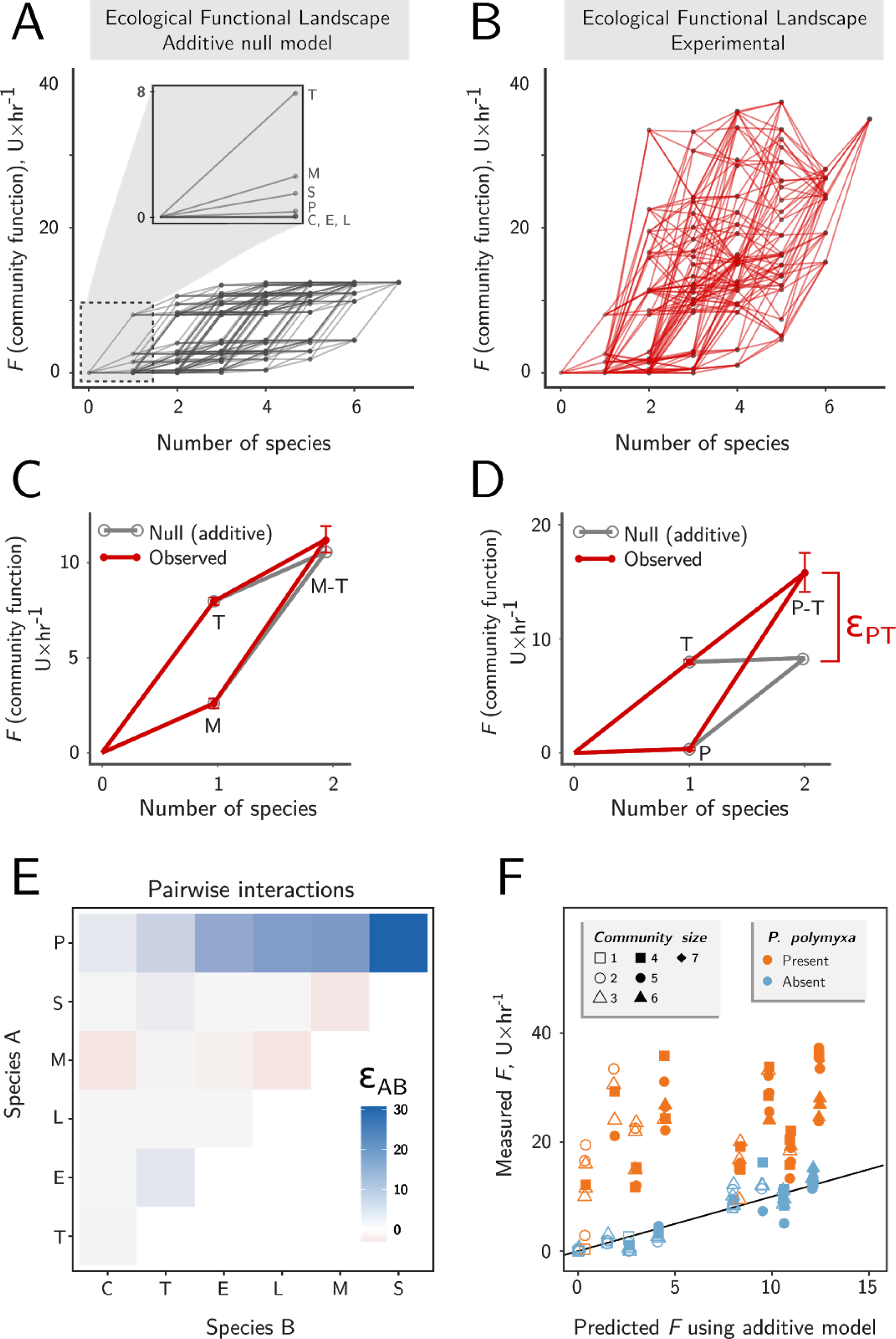
Pairwise interactions are ubiquitous in our microbial consortia. Cultures of the seven bacilli species grown at 30^°^C for 24 hrs were concentrated, washed and resuspended in growth media to assemble consortia containing one, two, three, four, five, six or seven species. Communities grew at 30^°^C for an additional 24hrs and the amylolytic rate of each supernatant was then measured (Methods). Species are designated as C, *B. cereus*; E, *B. megaterium*; L, *B. licheniformis*; M; *B. mojavensis*; P, *P. polymyxa*; S, *B. subtilis* and T, *B. thuringiensis*. **A.** Functional landscape expected from a pure additive biochemical model, as a function of species richness (*x* axis, richness measured as number of species in the consortia). The additive model assumes that the function (*F*; amylolytic rate) of the enzymes collectively secreted by the communities represents the sum of each individual species contribution (shown in the inset). **B.** Experimental functional landscape, where each point is the experimentally measured amylolytic function of a consortium. **C.** Example of a non-interacting pair (*B. mojavensis* and *B. thuringiensis*). Red lines show experimental measures (**±**SE), which are well predicted by the additive model (gray). **D.** Example of an interacting pair (*P. polymyxa* and *B. thuringiensis*) where the experimentally measured function of the pair (red) is not well predicted by the sum of the individual contributions (gray). The difference ε_PT_ quantifies the pairwise interaction. **E.** Pairwise interactions (ε_AB_) in all possible two-member species consortia. **F.** Comparison of the community function (amylolytic rate) experimentally measured as in Fig. 1D versus the function predicted by the additive model. Shape and color represent community size and presence of *P. polymyxa*, respectively.

To better understand these interactions and their implications, we quantified the deviations in the function of each consortium to the biochemical null in an approach resembling how epistasis quantifies gene-gene interactions in the context of fitness landscapes[6]. For instance in Fig. 2C-D we plot the Functional Landscape for two different pairwise consortia: one formed by *B. thuringiensis* and *B. mojavensis* (Fig. 2C) and a second one formed by *P. polymyxa* and *B. thuringiensis* (Fig. 2D). In gray, we show the expected function from the null model that assume that no interactions are present and predicts that the function of a pair of species *A* and *B (F*_*AB,*(Null)_) would be equal to the sum of the catalytic rates of each species in isolation (*V*_*A*_ and *V*_*B*_, so *F*_*AB,*(Null)_=*V*_*A*_*+V*_*B*_). On top of this prediction we plot in red the experimentally measured function of the pairwise consortia. As we see in Fig 2C, the null model approximates fairly well the function of the pairwise consortia formed by *B. thuringiensis* and *B. mojavensis*, suggesting that the pairwise interaction is weak for this pair. In contrast, the null model fails for the second consortia formed by *B. thuringiensis* and *P. polymyxa* (Fig. 2D), indicating the presence of a stronger functional interaction (which we denote by the symbol ε_AB_). The deviation between the null model prediction and the function of the consortia quantifies the pairwise functional interaction (i.e: *F*_*AB*_=*V*_*A*_*+V*_*B*_*+*ε_AB_).

The strength of pairwise functional interactions can be determined in the same manner for other pairs of species too, and the result is presented in Fig. 2E. This plot identifies one of the species, *P. polymyxa,* as the one having by far the strongest pairwise functional interactions. For pairs that do not involve *P. polymyxa*, pairwise interactions are weaker (and often negative) (Fig. 2E). Consistent with these findings, the additive “null” model is highly predictive of function (explaining 84.9% of the variance) for all of the consortia where *P. polymyxa* is absent, including three, four, five and six members (Fig. 2F). Yet, it grossly underestimates function for all of the consortia that contain *P. polymyxa*, irrespectively of community size (Fig. 2F).

### Quantifying high-order functional interactions in our simple consortia

Although pairwise interactions are in some cases sufficient to describe important effects in ecology, recent work has highlighted the need to incorporate high-order relations, focusing mainly on their role in population dynamics [25,33–35,48,49]. However, the role that high-order functional interactions may play in determining community function is not well understood, and has only recently began to be investigated [6,36]. Again borrowing from the study of complex genetic interactions[50], one could determine the contribution of high-order functional interactions to the community-level function by measuring how much the pairwise functional interactions depend on their ecological context (e.g. which other members of the community are present) (Fig. 3A). To determine to which extent pairwise interactions depend on the presence of other species, we measured the amylolytic rate for every possible pairwise and three-member consortia, and then followed the approach outlined in Fig. 3A to determine how the presence of a third species altered the strength of a pairwise interaction. The results are summarized in Fig. 3B, which exhibits a strong trend: the presence of *P. polymyxa* substantially alters the pairwise functional interactions between every two other species, and the pairwise interactions that include *P. polymyxa* also exhibit substantial changes when a third species is present.

**Figure 3.**
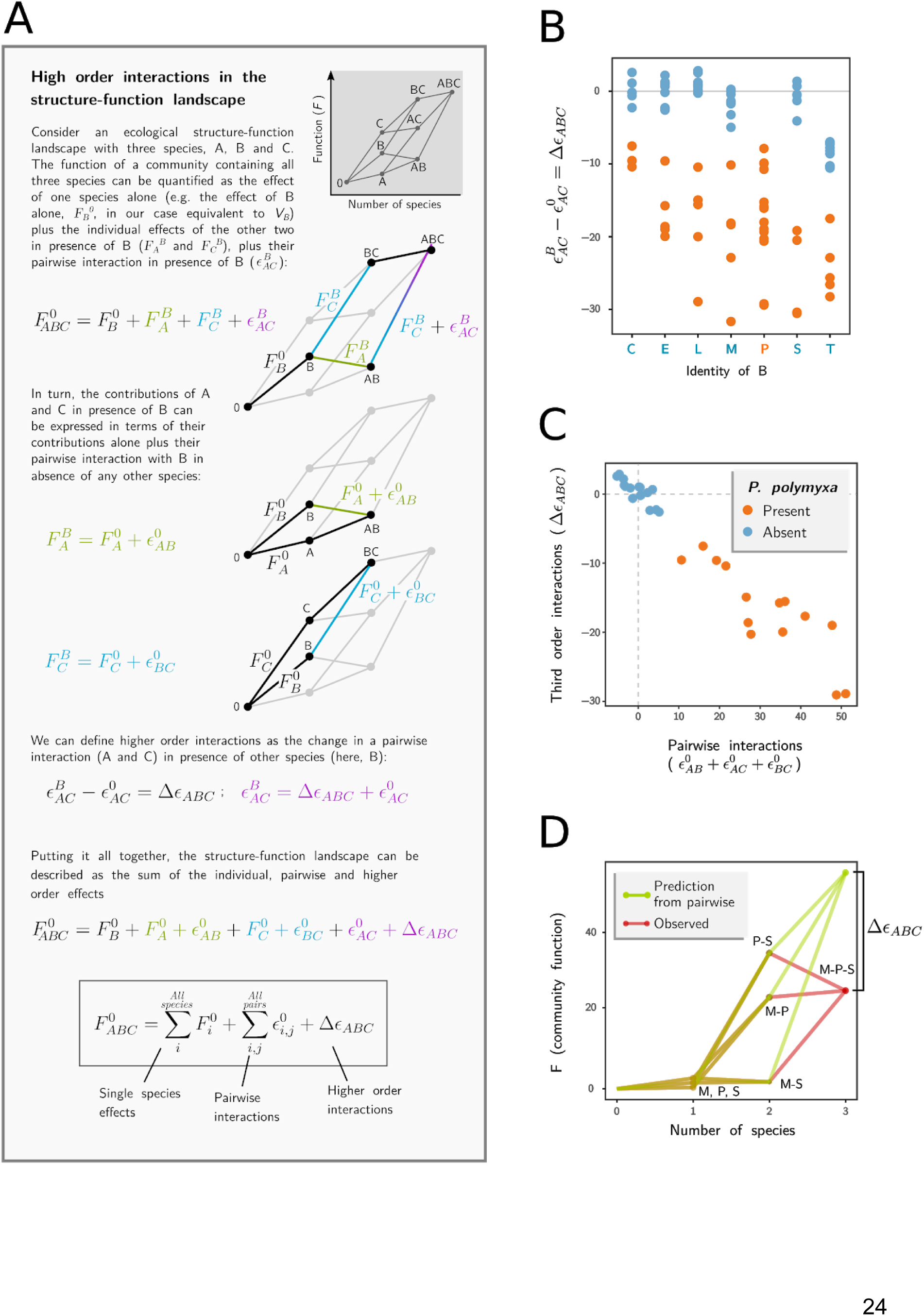
Higher order interactions in simple amylolytic consortia. **A.** Box explaining how single, pairwise and higher order interactions in the function of microbial consortia can be separated and quantified using Ecological Functional Landscapes. **B.** Third order interactions in trios 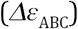, quantified as the difference in the interaction when a pair “AC” is grown alone 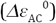versus in the presence of a third species “B” 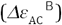 whose identity is shown in *x* axis. Species are designated as in Fig. 2. Trios containing *P. polymyxa* are shown in orange, and in blue otherwise. **C.** Third-order interactions (y axis) are strongly anti-correlated to pairwise interactions (x axis) Pearson’s 𝜚=-0.96, P<10^−10^. **D.** Quantification of a third-order interaction between *P. polymyxa* (P), *B. mojavensis* (M) and *B. subtilis* (S) suggests redundancy between *B. mojavensis* and *B subtilis* when *P. polymyxa* is present.

This analysis shows that third-order functional interactions are present in our consortia, and they are particularly strong when they involve *P. polymyxa*. As a result, a model that incorporates the sum of all single species contributions and pairwise interactions for a three-member consortia, vastly overestimates the amylolytic function of our communities. An example is given in Fig. 3C, which shows a three member community formed by *P. polymyxa, B. cereus* and *B. megaterium*. None of the three have a strong amylolytic activity in monoculture, but the two pairs that include *P. polymyxa* have high amylolytic rates, driven by strong pairwise interactions. However, as shown in Fig. 3D, the trio has much lower function than would be expected by adding together every possible pairwise interaction. Third-order interactions (but not first order interactions; Fig S2) exhibit a strong negative correlation with the sum of all pairwise interactions (Pearson’s ρ=-0.96, P<10^−10^) (Fig. 3D), suggesting that pairwise interactions do not add up as one may predict based on the Functional Landscape model in Fig. 3A. Instead, the result in Fig. 3D shows that pairwise interactions combine sub-additively, indicating that pairwise interactions in communities that contain *P. polymyxa* may be redundant.

### Redundancy of ecological interactions in multi-species consortia

The results reported in Fig. 3D indicate that in multi-species consortia, the pairwise interactions involving *P. polymyxa* are functionally redundant. The strain of *P. polymyxa* we used in our consortia is a biotin auxotroph [51], and it grows poorly in our growth medium, which does not include a vitamin supplement (Fig. S3). This led us to hypothesize that the positive pairwise interactions between every other community member and *P. polymyxa* may arise as a result of facilitation towards *P. polymyxa* (Fig. 4A). To test this hypothesis, we determined the number of Colony Forming Units of *P. polymyxa* in monoculture as well as in co-culture with each of the other members of the consortia. Consistent with our hypothesis we find that *P. polymyxa* is substantially facilitated by every other member of the consortium (Fig. 4B), but their facilitative effects are redundant: co-culturing *P. polymyxa* with three or more members of the consortium produces the same growth stimulation than that observed in pairwise co-culture (Fig. 4C).

**Figure 4.**
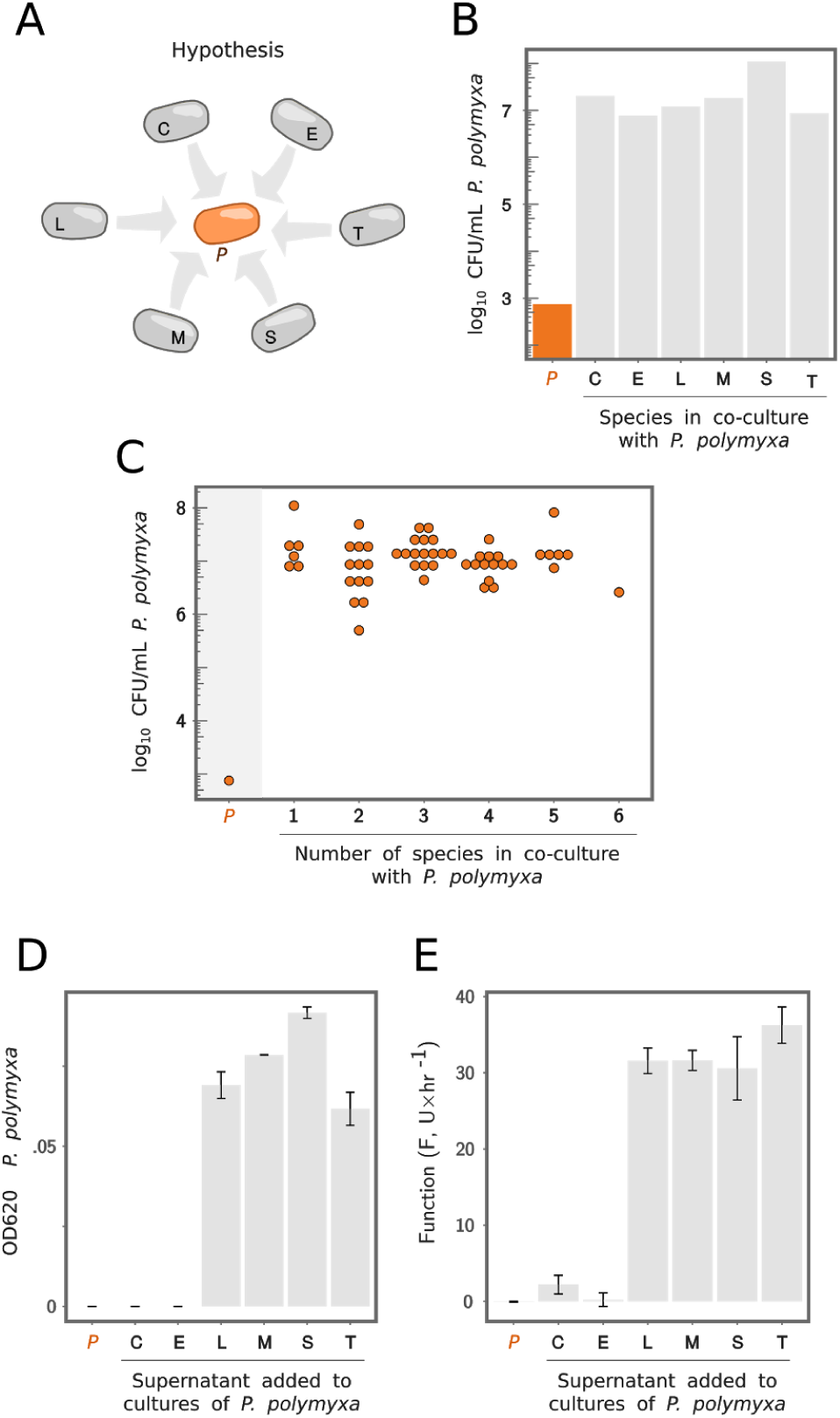
Redundancy in the ecological facilitation of *P. polymyxa* explains higher-order interactions. **A.** If *P. polymyxa* growth can be facilitated by any other species, this facilitation could be redundant. Species are designated as in Fig. 2. **B.** *P. polymyxa* grows in the presence of any other species (gray bars), but not in monoculture (orange bar). CFU/mL was obtained by colony counting of serially diluted cultures after 24hrs of growth at 30^°^C. **C.** *P. polymyxa* grows to a comparable density in the presence of concurrent species, irrespective of their number (x axis). This strongly suggests redundancy in the facilitation mechanism. **D.** In contrast to monoculture in minimal medium (orange), *P. polymyxa* exhibits growth when cultured in media supplemented with the filtered supernatant of other species (1:10 dilution, Methods). **E.** Function (amylolytic activity, U × hr^−1^) of *P. polymyxa* cultured as in panel D.

Growth stimulation is also observed when *P. polymyxa* is grown in media supplemented with the filtered supernatant of every other species in monoculture (with the exception of *B. megaterium* and *B. cereus*, Fig. 4D). This ecological facilitation also results in a marked increase in the amylolytic activity of *P. polymyxa* (Fig. 4E). To ensure that this increase in amylolytic activity is not due to the activity of extracellular amylases carried over in the *Bacillus* filtrates, we filtered these a second time through a 30 KDa filter membrane (Methods), which is small enough to let vitamins such as biotin through (MW∼0.244 KDa) but blocks the passage of amylases (MW >50 KDa). Indeed, the filtrate through the second 30 KDa filter exhibited no amylolytic activity, demonstrating that amylases are not present.

These results suggest that cross-feeding is sufficient to stimulate the growth of *P. polymyxa*, which also leads to a larger amylolytic activity (Fig. 4E), comparable to the highest levels observed among communities (see Fig. 2B). We also find that all of the *Bacillus* species in our consortia are able to facilitate the growth of *P. polymyxa*, supporting our hypothesis that this facilitation is redundant. In other words, any combination of the *Bacillus* species stimulates the growth of *P. polymyxa* growth about as much as any of these species do separately (Fig. 4C and Fig. S5). Because all other members of the consortium similarly stimulate the growth of *P. polymyxa*, and their combined effects are less than additive, high-order corrections are strongly negative and scale with the overestimation produced by adding together the second-order pairwise functional interactions, which explains the strong correlation between third-order and second order interactions reported in Fig. 3C.

### The strength of high-order interactions increases as the communities grow in size

The importance of high-order interactions is reflected by the poor predictive power of the pairwise model (Fig. 5A) for consortia with three or more species. This last result prompted us to ask whether the importance of high-order interactions would grow as communities increase in size. As shown in Fig. 5B, the ability of the pairwise to predict community function declines as more and more species are added to the consortia. As shown in Fig. S4, the same is true even if we include third-order interactions into the model.

**Figure 5.**
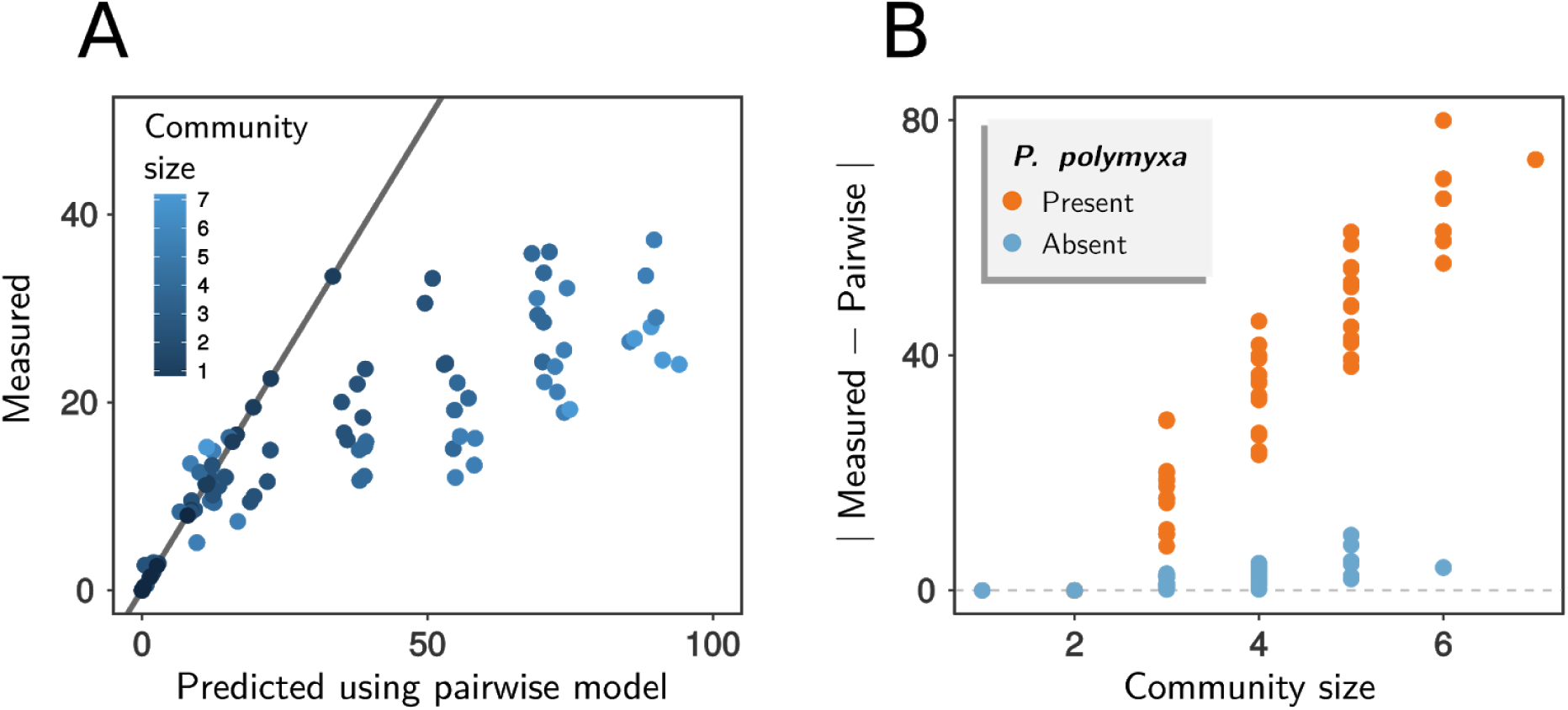
The strength of high-order interactions increases with community size. **A.** Plot showing the prediction from the pairwise model against the experimentally measured value for communities of different size (represented by point color). **B.** Strength of higher order interactions, measured as the absolute value of the difference between the function determined experimentally(amylolytic activity, U × hr^−1^) and the expected one using the model including only single and pairwise effects.

## Discussion

Functional interactions are observed when the contribution of a community member, or set of members, to a community-level function depends on the presence or absence of other species. For instance, pairwise interactions reflect how the contribution of a single species to a community function (e.g. in the case explored above, the amylolytic activity of the enzymes it secretes) depends on whether it is alone or in co-culture with a second species. Likewise, third order interactions capture how the function of a pair of species, (e.g. the amylolytic rate of the enzymes secreted by the pair) is altered when a third species is present. This simple idea, based on the study of fitness landscapes and complex interactions in genetics, allowed us to decompose the function of a community into the contributions of single species and the interactions that modulate these contributions, and can be used to shed light onto the role played by high order interactions in community function. As we have shown here, and others have shown before in different contexts [37], a null model of how the functional contributions of multiple species should combine to determine the community function is essential in order to unequivocally identify interactions through this approach.

Most community-level functions of interest are highly complex. For instance, in consortia designed for biotransformation, multiple chemical steps (often carried out by different species) are typically required to convert waste materials into high-value products [8,9]. For host associated communities, functions of interest typically include effects that the community has on its host, be it on its health [15,52,53], behavior [5], or life history traits [3,6]. Thus, these functions involve not only biochemical processes carried out by the community, but the host’s response to them. Given this complexity, it is not obvious how functional contributions from many species should combine together to determine the community function.

A similar problem has been found in the study of complex genetics. Most models in genetics assume that the effects of non-interacting mutations are additive (e.g. [41,42,54]). Although additivity can be used for lack of a better alternative, many functions in complex biological and biochemical systems are intrinsically non-additive even in the absence of direct interactions [44], potentially leading to misinterpretations of what are and are not interactions. This is well known in genetics, and it has lead to the argument that pairwise or higher order interactions may be statistical artifacts in the presence of non-linearities [37]. Unfortunately, what the form of the null model should be is far from obvious when the function or phenotype of interest is complex and difficult to model mechanistically. Some encouragement comes from the recent application of similar methods to the study of interactions between antibiotics [46] and cancer drugs [47]. The combinatorial effect of multiple drugs could be predicted in a model-free manner from single drug effects and pairwise interactions alone, suggesting that low dimensional representations and a null additive model may be predictive even for highly complex biological processes for which a detailed mechanistic interpretation is not available [46,47].

Our approach here has been to examine a function for which a quantitative biochemical model could be derived, thus offering a direct estimate of interactions that does not rely on a priori assumptions. By focusing on a simple, tractable model system for which we have a theoretically grounded mechanistic null model, we have been able to unambiguously determine the presence of pairwise and higher interactions in our model consortia. Our results emphasize the importance of high-order interactions in determining community-level functions, even for a decidedly non-complex function: one that could be independently carried out by each species in isolation, and that does not require more than a single gene in each species.

One potential feature of our study is worth discussing. Our null model predicts that the amylolytic rate of the enzymes secreted by a multi-species consortium should be equal to the sum of the amylolytic rates of the enzymes released by each species in monoculture. One could argue that, when grown in co-culture in the same bioreactor, each species would have access to less resources than they do in monoculture, as they are competing for the same pool of nutrients. It might thus seem implausible that these species would ever combine additively, as the density that each species reaches in co-culture would be lower than it is in isolation. We note, however, that competition is a form of ecological interaction that could have effects on function, and may very good lead to the function of a consortia being different than additive. In spite of this, we find that consortia that lack *P. polymyxa* are actually quite close to the additive functional prediction, (see Fig. 3D) even though competitive interactions are frequent in these consortia (Fig. S5). Moreover, the largest deviation we see from additivity come from consortia that contain *P. polymyxa*, and in those cases the deviation is always positive (the function of these consortia is always larger than the predicted by the additive model), and this is mediated by facilitation, rather than competition.

It is yet unclear what these findings mean for the complex functions that are of interest in most natural and synthetic systems. By comparison to an additive null model, high-order interactions have been reported in a set of recent experiments where the microbiome of *D. melanogaster* was combinatorially reconstituted to map its effects on several life history traits of the fly, such as its longevity or developmental time [6]. The confluence of findings between this study and ours suggest that high-order functional interactions may be generically prevalent in microbial communities, presenting a fundamental challenge to predict their function from the bottom-up. While this may be seen as a disappointment, it is not unusual in ecology, nor in complex systems in general, that the whole is more than the sum of its parts. Understanding exactly how this complexity works and how the parts come together to produce complex quantitative traits has lead to fruitful research in other fields, from genetics to metabolism to neuroscience. We hope that our findings will help motivate similar efforts in microbial ecology.

## Methods

### Strains and Media

Bacterial strains were obtained from ATCC (Manassas, VA) with the following designations: *Bacillus subtilis* (ATCC 23857), *Bacillus megaterium* (ATCC 14581), *Bacillus mojavensis* (ATCC 51516), *Paenibacillus polymyxa* (ATCC 842), *Bacillus thuringiensis* (ATCC 10792), *Bacillus licheniformis* (ATCC 14580). *Bacillus cereus* was isolated from a soil sample in Boston, Massachusetts. Cell stocks were prepared according to manufacturer instructions and stored at −80°C in 40% glycerol.

Basic growth minimal media (1x bSAM) was prepared from 10x concentrated stocks of bM9 Salts containing Na_2_HPO_4_ × 2H_2_O (85.4 g/L; Bioworld), KH_2_PO_4_ (30 g/L; Fisher Scientific), NaCl (5 g/L; Sigma Aldrich), NH_4_Cl (10 g/L; Fisher Scientific), supplemented with 0.04% synthetic complete amino acids (w/v; Sunrise Science Products), 0.1% starch (w/v; soluble, Sigma Aldrich), 1% trace mineral supplement (v/v; ATCC), CaCl_2_ (0.1mM; Sigma Aldrich), and MgSO_4_ (2mM; Fisher Scientific). Starch assay media consisted of 2x bM9 salts, supplemented with 0.2% starch (w/v), CaCl_2_ (0.2mM) and MgSO_4_ (4mM).

### Culture inoculation and combinatorial assembly

Strains were streaked out on BE Starch Agar (0.3% beef extract, 1% starch) plates and grown for 24 hours at 30^°^C. Seed cultures were started from several colonies (depending on colony size), inoculated into three mL 1x bSAM and grown still at 30^°^C for 24 hours. Cultures were then transferred to a 96 well plate (Corning Cat. No. 3596) and the optical density (620 nm) of 100 µL was measured (Multiskan Spectrophotometer; Fisher Scientific). Cells were harvested by centrifugation at 3500 rpm for 15 min, washed twice with 1x phosphate buffered saline and suspended in fresh 1x bSAM media at a concentration of 5×10^5^ cfu/µL. Monocultures or combinatorially assembled communities of the seven bacilli species were prepared by inoculating two µL from each seed culture into 96 deep-well plates (VWR) containing 500 µL of 1x bSAM. Plates were covered with Aerogel film (VWR) and incubated still at 30^°^C for another 24hrs. Optical density (620 nm) of the grown cultures was measured as above at the end of the new incubation period.

### Quantifying growth of P. polymyxa in pairwise co-cultures

Combinatorially assembled communities were incubated at 30°C for 24hrs and then stored at −80°C in 40% glycerol. To measure the amount of *P. polymyxa* colony forming units (cfu) in each mixture, ∼20µl of the frozen stock was melted and serially diluted 1:10 up to 1:10^5^. 50µl of dilutions 1:10^2^, 1:10^3^, 1:10^4^ and 1:10^5^ were then plated onto BE Starch and incubated at 30^°^C for 48hrs. Plates were scanned with an EPSON Perfection V700-V750 scanner at a 300dpi resolution and cfu/mL were recorded with the ImageJ plugin Cell Counter. Colony morphology was different enough to easily allow differential identification of *P. polymyxa* from all other species.

### Cross-feeding Assays

The seven bacilli strains were individually grown in three mL of 1x bSAM at 30^°^C for 24 hrs. To obtain the the supernatants for cross-feeding assays, cultures were pelleted at 3500 rpm for 15 min and the supernatants were carefully transferred to clean sterile tubes and kept at room temperature. Cells were processed as described above. Supernatants were then transferred to 0.2 µm spin filter columns (VWR 82031-358) and centrifuged at 14000 x*g* for 5 minutes. The flow-through was subjected to a second filtration step through a 30 KDa centrifugal device (Nanosep 3K; Pall OD003C33) at 5000 x*g* for 10 min. 50 µL of undiluted supernatants were mixed with 250 µL 2x bSAM and 200µL water in 96 deep-well plates. Two µL containing 10^6^ cfu of *P. polymyxa* cells were inoculated into this media and grown still at 30^°^C for 24 hours. Prior to filtering, all devices were sanitized with 70% ethanol, according to manufacturer instructions.

### Determination of amylolytic rates

Starch hydrolysis assays were based on a Lugol iodine staining method for starch [55]. Lugol Staining solution was prepared with 390 mL water: 60 mL Lugol’s Iodine stain (Sigma Aldrich). Supernatants containing extracellular amylases were prepared for enzymatic assays by applying 330µL of homogeneously suspended bacterial cultures grown for 24 hrs directly to a 96 well 0.2µm Acroprep filter plate (Pall Corp., Cat. No. 8019) fitted to a 96 well collection plate (Corning, Cat. No. 3596) with the metal collar adaptor (Pall Corp., Cat. No. 5225) and centrifuged for 20 minutes at 1500 xg in a table top centrifuge (Eppendorf 5810).

Assays, in 96 well plates, contained varying volumes of filtered supernatant (100, 50, 25, 10 and 5µL), 100µL 2x SAM and water to a total reaction volume of 200 µL. Control reaction plates were also prepared for either no enzyme (100µL 2x SAM and 100µL water) or no starch with supernatant (100µL water and 100µL supernatant). Reactions, assembled with pre-warmed SAM, were incubated at 30^°^C to the desired time point, and quenched by transferring 50 µL to a solution containing Lugol Iodine stain (130 µL water and 20 µL Lugol per well). Starch degradation was quantified by measuring the optical density at 690 nm.

### Michaelis-Menten kinetic model of starch hydrolysis

Our model is based on the theory of chemical kinetics [56], which predicts that when a reaction can occur independently and in parallel through *n*_*S*_ distinct channels, the overall rate of the reaction is the sum of the rates for each of the independent channels [57]. We here consider that starch is broken down in a two-step Michaelis-Menten scheme, with both steps having equal rate *V* _*j*_ = *k*_*cat, j*_ [*E*]_*j*_ where *j* = 1, 2, …, *n*_*S*_. We make the further assumption here that substrate concentration [*S*] is much larger than the Michaelis constant *K*_*M*_, so that the enzyme is always saturated. Previous characterization of *K*_*M*_ for various bacilli amylases acting on non-soluble amylases are in the range of ∼1-5 mg/mL [58,59], indicating that this assumption is reasonable. Each step is catalyzed by the *n*_*S*_ types of enzymes released by the *n*_*S*_ species in the consortia. Since all enzymes break down starch independently from each other, the velocity of each step is thus:

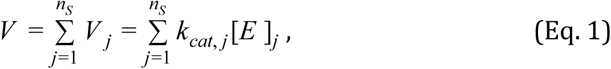

and therefore the fraction of starch degraded by the enzymes after a time *t* is:

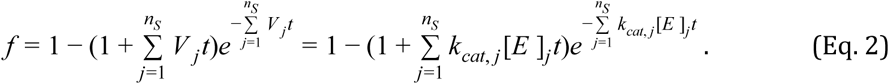

Equation [4] gives us the fraction of starch degraded over a time *t* by a cocktail of enzymes of rates ^*V*^ *j* ^= *k*^*cat, j* ^[*E*]^*j* that act independently on the substrate, and whose catalytic rates are not affected by each other’s presence.

A fundamental prediction of this model is that there exists a symmetry between the enzyme concentration ([*E*]) and the incubation time (*t*). In other words, the same amount of starch would be degraded by increasing the incubation time 10-fold and leaving the enzyme concentration constant, as it would by leaving the incubation time constant and increasing the enzyme concentration by 10-fold. This prediction was proven to be valid for a wide range of values of time (from 10min to 120min) and enzyme concentration (0.0078mg/mL to 0.125mg/mL) (Fig. S1).

### Determining pairwise and third order interactions

High-order interactions can be quantified by following the scheme in Fig. 3A, and briefly summarized below: Denoting by 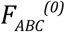 the starch hydrolysis rate of the enzymes secreted by a three-species community, we can decompose this function in terms of the functional contributions from each of the single species in isolation 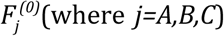(where *j=A,B,C*), and which in this case are equal to the catalytic rates of the enzymes secreted by each species in monoculture 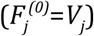; the pairwise interactions measured from each pair in isolation (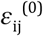, where *i,j=A,B,C*), and a term that captures the high-order three-way interactions when the three species are together in co-culture 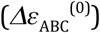:

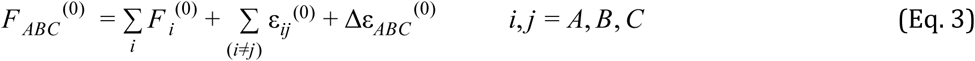

Since 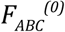 is the net function of the three-species community, which can be measured experimentally; and all of the 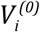 and 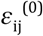 parameters can also be measured independently from the function of the monocultures and pairwise co-cultures, we can experimentally determine the strength of three-way interactions 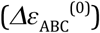 as:

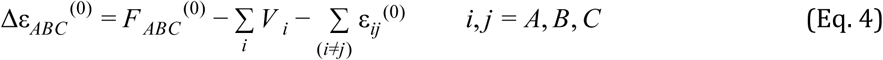

### Determining fourth and higher-order interactions

For communities of size *N* (larger than three-species), Equation 3 can be expanded as

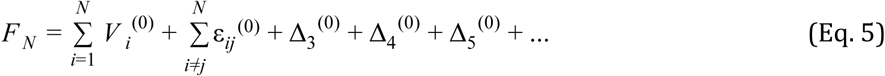

Where the collection of high order interactions can be defined as the sum of three-way 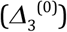 four-way 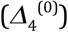, five-way 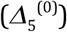 interactions in isolation, etc., and can be determined experimentally as:

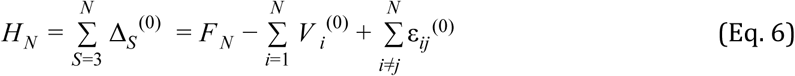

## Acknowledgements

We wish to express our gratitude to the Sanchez laboratory at Yale and to the Poyatos lab at the CNB-CSIC for helpful discussions.

## Funding

The funding for this work partly results from a Scialog Program sponsored jointly by Research Corporation for Science Advancement and the Gordon and Betty Moore Foundation through grants to Yale University (A.S.). This work was also supported by a young investigator award from the Human Frontier Science Program to A.S. (RGY0077/2016) and by grant FIS2016-78781-R from the Spanish Ministerio de Economía y Competitividad (J.F.P.).

## Author Contributions

J.F.P. and A.S. conceived the project, designed the experiments, and provided guidance; A.S-G., J.F.P., M.L.O. and A.S. performed experiments; A.S-G., D.B., M.L.O., J.F.P. and A.S. performed data analysis; A.S-G., D.B., J.F.P. and A.S. discussed the results; A.S-G., D.B., M.L.O.,J.P. and A.S. wrote the paper.

## Competing Interests

The authors declare that no competing interests exist in relation to this manuscript.

## Supplementary information

**Figure S1.**
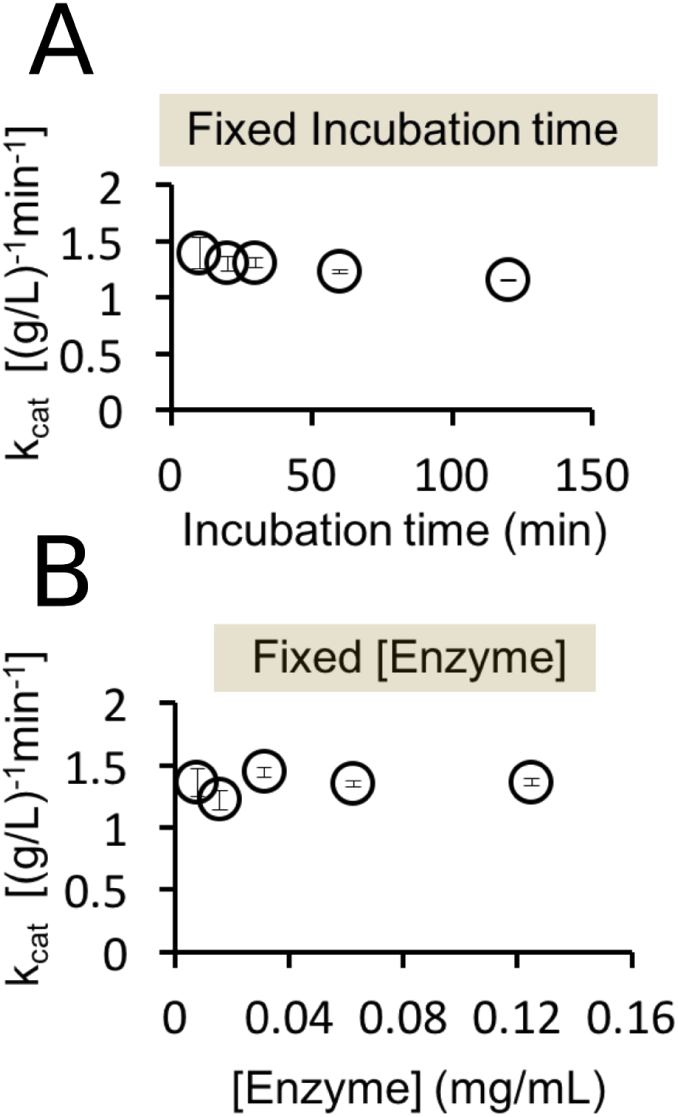
The model predicts an invariance between the time of incubation and the enzyme concentration. This was confirmed experimentally by either **A.** fixing the incubation time *t* and changing the enzyme concentration *[E]* or **B.** fixing the enzyme concentration and changing the incubation time. The fitting parameter *k*_*cat*_ obtained is consistent across both conditions and over several orders of magnitude for both *[E]* and *t*, indicating that the two-step Michaelis-Menten model is a reasonable quantitative mechanism of starch hydrolysis by extracellular amylases.

**Figure S2.**
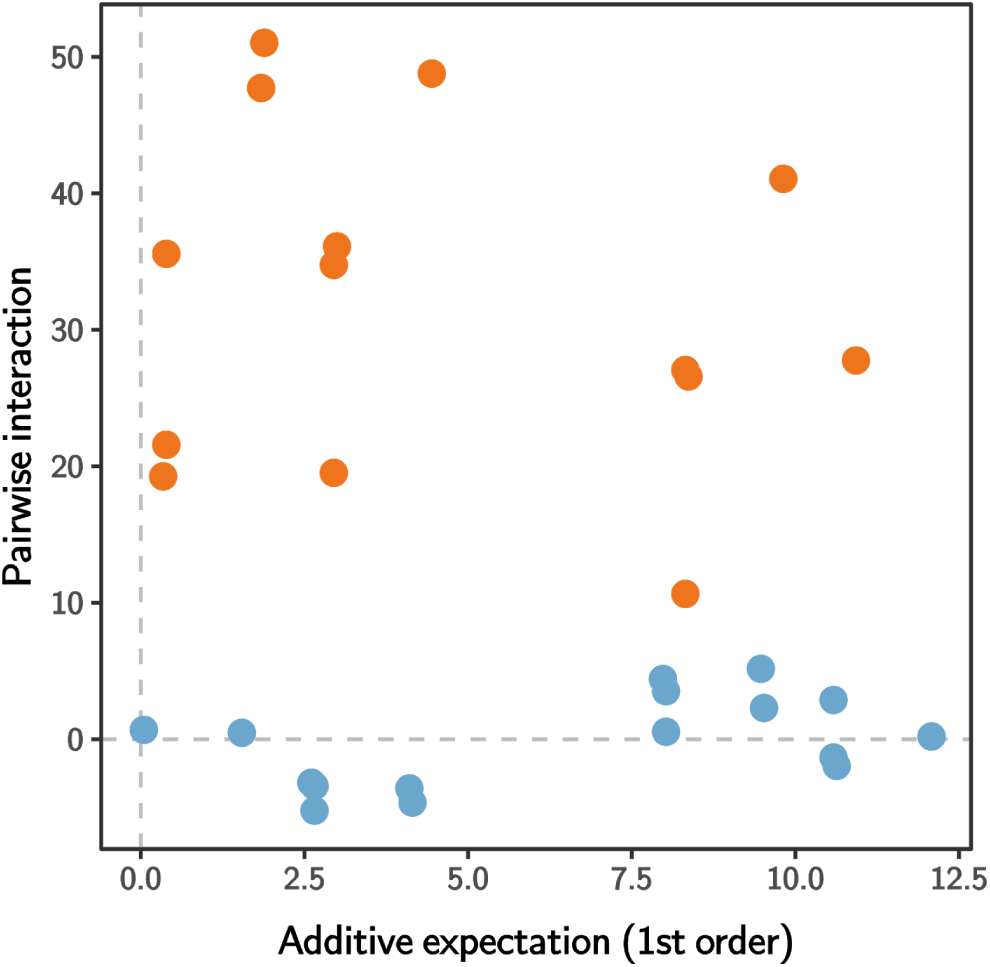
Sum of pairwise interactions is not correlated to additive expectation in three-species communities. Groups containing *P. polymyxa* are shown in orange, and in blue otherwise.

**Figure S3.**
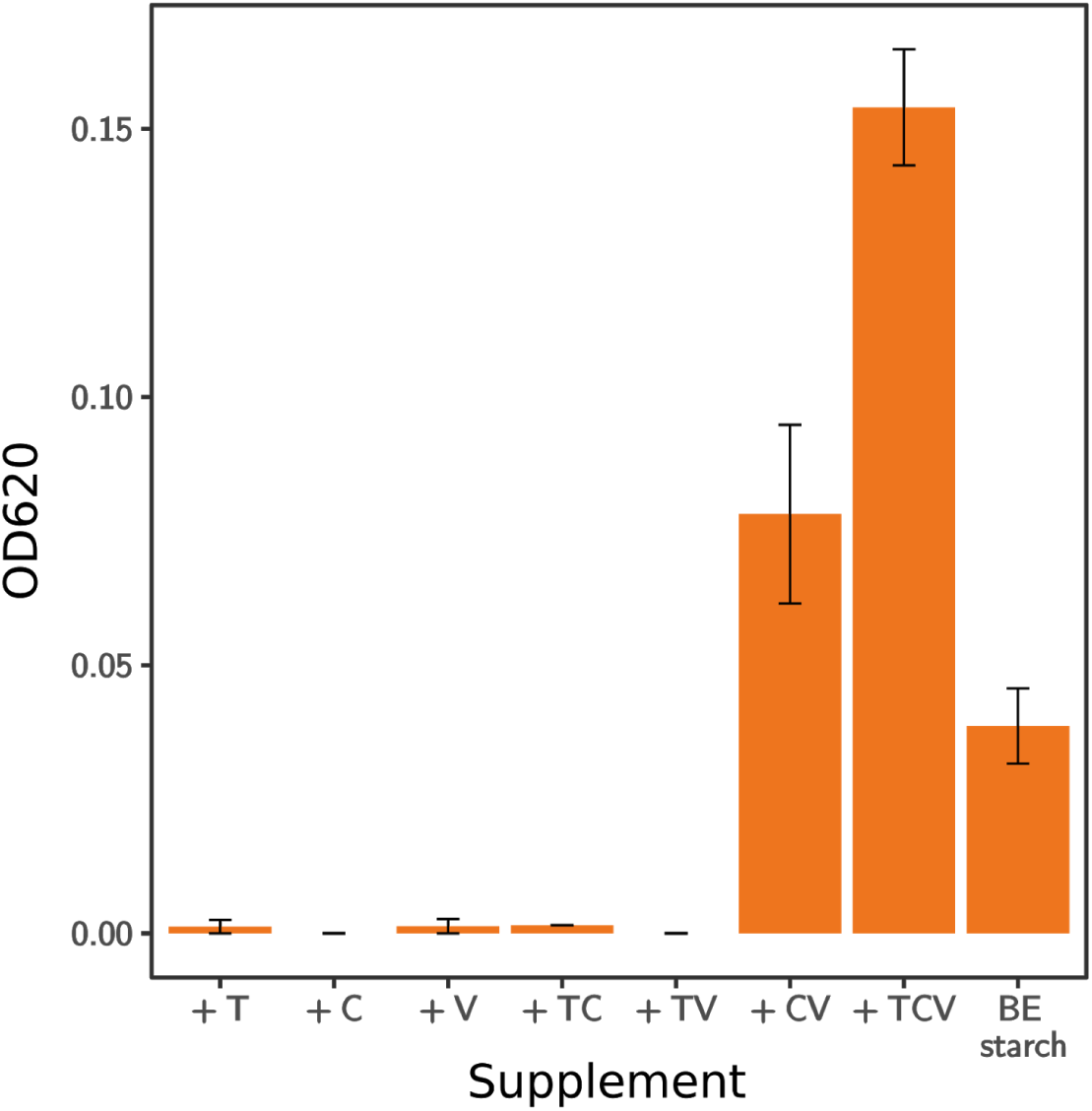
*P. polymyxa* grows better in media supplemented with vitamins. Optical density (620nm) of cultures of *P. polymyxa* grown at 30^°^C for 24 hrs in rich media (BE starch) or 1x bSAM supplemented, as indicated, with a combination of 1% trace mineral supplement (T); 0.04% synthetic complete amino acids (C) or 1% vitamin supplement (V; v/v, ATCC). Growth of *P. polymyxa* in media containing the three supplements (TCV) shows a marked increase, whereas the media routinely used in this work (TC) poorly sustains its growth. Plots show the media for two replicates and the standard error.

**Figure S4.**
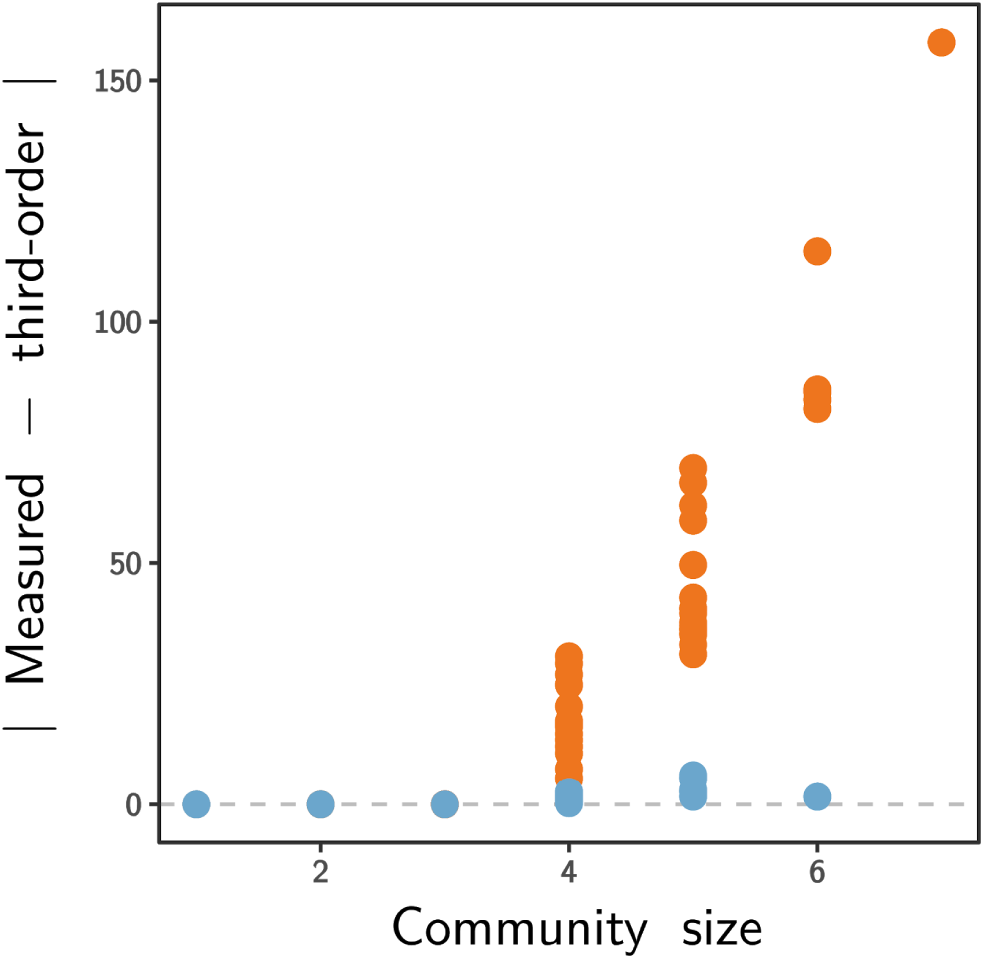
Strength of higher order interactions, measured as the absolute value of the difference between the measured function and the expected one using the model including only single, pairwise and third-order effects. Orange and blue dots represent communities with or without *P. polymyxa*, respectively.

**Figure S5.**
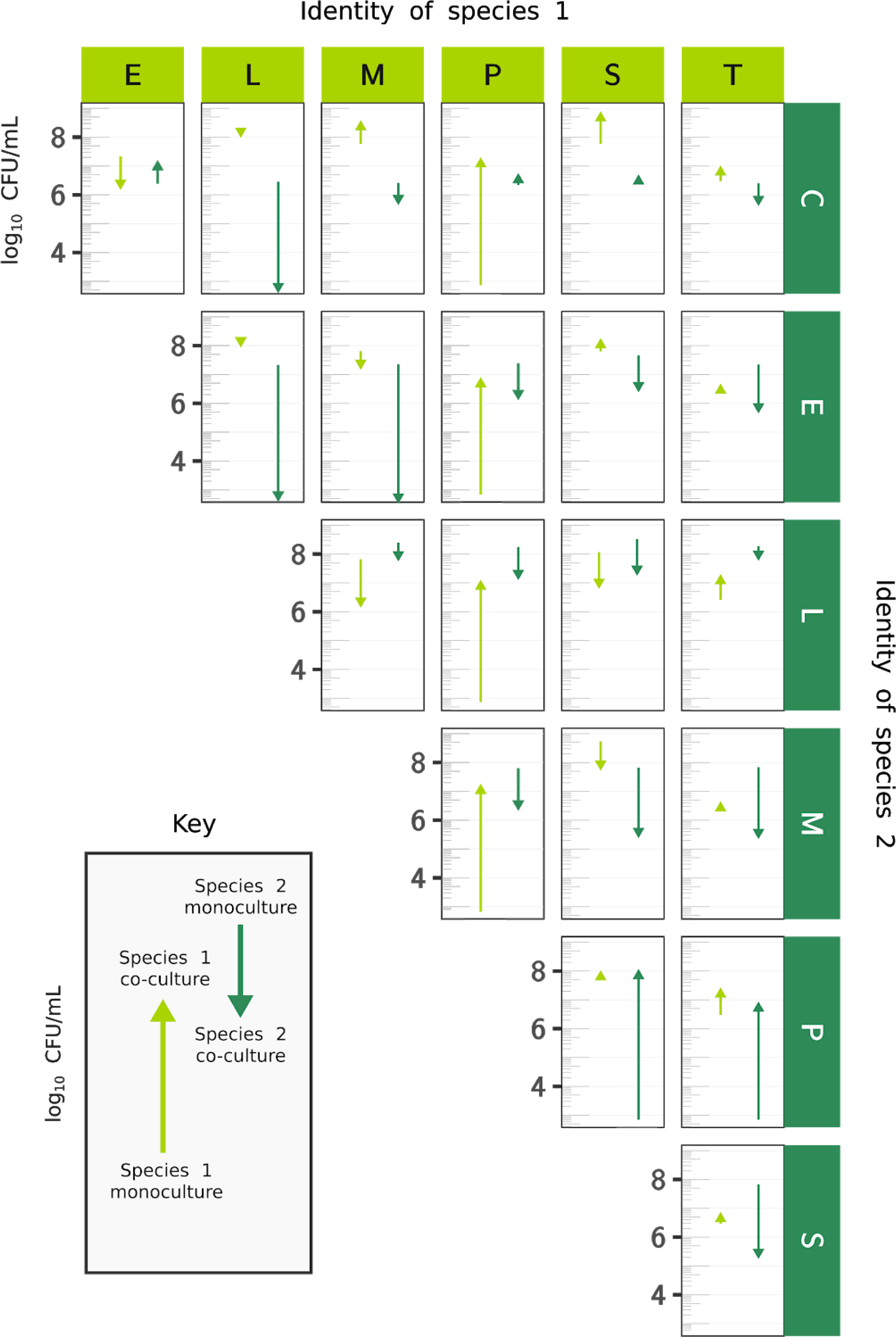
Changes in CFU/mL observed after co-culturing the different bacilli species in pairs. Cultures of the seven bacilli species grown at 30^°^C for 24hrs were concentrated, washed, resuspended in growth media to inoculate monocultures or pairs and were grown at 30^°^C for an additional 24hrs. Species (light and dark green) are designated as C, *B. cereus*; E, *B. megaterium*; L,*licheniformis*; M; *B. mojavensis*; P, *P. polymyxa*; S, *B. subtilis* and T, *B. thuringiensis*. Upward arrows denote an increase in CFU/mL from the value obtained when the two species are grown in monocultures to the value obtained after the same two species were co-cultured for 24hrs. Downward arrows indicate a decrease in CFU/mL.

## References

1. Thompson LR, Sanders JG, McDonald D, Amir A, Ladau J, Locey KJ, et al. A communal catalogue reveals Earth’s multiscale microbial diversity. Nature. 2017;551: 457–463.

2. Falkowski PG, Fenchel T, Delong EF. The microbial engines that drive Earth’s biogeochemical cycles. Science. 2008;320: 1034–1039.

3. Panke-Buisse K, Lee S, Kao-Kniffin J. Cultivated Sub-Populations of Soil Microbiomes Retain Early Flowering Plant Trait. Microb Ecol. 2016; doi:10.1007/s00248-016-0846-1

4. Wagner MR, Lundberg DS, Coleman-Derr D, Tringe SG, Dangl JL, Mitchell-Olds T. Natural soil microbes alter flowering phenology and the intensity of selection on flowering time in a wild Arabidopsis relative. Ecol Lett. 2014;17: 717–726.

5. Leitão-Gonçalves R, Carvalho-Santos Z, Francisco AP, Fioreze GT, Anjos M, Baltazar C, et al. Commensal bacteria and essential amino acids control food choice behavior and reproduction. PLoS Biol. 2017;15: e2000862.

6. Gould AL, Zhang V, Lamberti L, Jones EW, Obadia B, Gavryushkin A, et al. High-dimensional microbiome interactions shape host fitness [Internet]. bioRxiv. 2017. p. 232959. doi:10.1101/232959

7. Brenner K, You L, Arnold FH. Engineering microbial consortia: a new frontier in synthetic biology. Trends Biotechnol. 2008;26: 483–489.

8. Minty JJ, Singer ME, Scholz SA, Bae C-H, Ahn J-H, Foster CE, et al. Design and characterization of synthetic fungal-bacterial consortia for direct production of isobutanol from cellulosic biomass. Proc Natl Acad Sci U S A. 2013;110: 14592–14597.

9. Shong J, Jimenez Diaz MR, Collins CH. Towards synthetic microbial consortia for bioprocessing. Curr Opin Biotechnol. 2012;23: 798–802.

10. Lindemann SR, Bernstein HC, Song H-S, Fredrickson JK, Fields MW, Shou W, et al. Engineering microbial consortia for controllable outputs. ISME J. 2016;10: 2077–2084.

11. Foo JL, Ling H, Lee YS, Chang MW. Microbiome engineering: Current applications and its future. Biotechnol J. 2017;12. doi:10.1002/biot.201600099

12. Jousset A, Eisenhauer N, Merker M, Mouquet N, Scheu S. High functional diversity stimulates diversification in experimental microbial communities. Sci Adv. 2016;2: e1600124.

13. Hu J, Wei Z, Friman V-P, Gu S-H, Wang X-F, Eisenhauer N, et al. Probiotic Diversity Enhances Rhizosphere Microbiome Function and Plant Disease Suppression. MBio. 2016;7. doi:10.1128/mBio.01790-16

14. Mueller UG, Juenger T, Kardish M, Carlson A, Burns K, Smith C, et al. Artificial Microbiome-Selection to Engineer Microbiomes That Confer Salt-Tolerance to Plants [Internet]. bioRxiv. 2016. p. 081521. doi:10.1101/081521

15. Mueller UG, Sachs JL. Engineering Microbiomes to Improve Plant and Animal Health. Trends Microbiol. 2015;23: 606–617.

16. Blouin M, Karimi B, Mathieu J, Lerch TZ. Levels and limits in artificial selection of communities. Ecol Lett. 2015;18: 1040–1048.

17. Widder S, Allen RJ, Pfeiffer T, Curtis TP, Wiuf C, Sloan WT, et al. Challenges in microbial ecology: building predictive understanding of community function and dynamics. ISME J. 2016; doi:10.1038/ismej.2016.45

18. Vorholt JA, Vogel C, Carlström CI, Müller DB. Establishing Causality: Opportunities of Synthetic Communities for Plant Microbiome Research. Cell Host Microbe. 2017;22: 142–155.

19. Konopka A, Lindemann S, Fredrickson J. Dynamics in microbial communities: unraveling mechanisms to identify principles. ISME J. 2015;9: 1488–1495.

20. Brenner K, You L, Arnold FH. Response to Goldman and Brown: Making sense of microbial consortia using ecology and evolution. Trends Biotechnol. Elsevier; 2009;27: 4.

21. Goldman RP, Brown SP. Making sense of microbial consortia using ecology and evolution. Trends Biotechnol. 2009;27: 3–4, author reply 4.

22. Momeni B, Xie L, Shou W. Lotka-Volterra pairwise modeling fails to capture diverse pairwise microbial interactions. Elife. 2017;6. doi:10.7554/eLife.25051

23. Tikhonov M. Community-level cohesion without cooperation. Elife. 2016;5. doi10.7554/eLife.15747

24. Lu N, Sanchez-Gorostiaga A, Tikhonov M, Sanchez A. Cohesiveness in microbial community coalescence [Internet]. bioRxiv. 2018. p. 282723. doi:10.1101/282723

25. Goldford JE, Lu N, Bajic D, Estrela S, Tikhonov M, Sanchez-Gorostiaga A, et al. Emergent Simplicity in Microbial Community Assembly [Internet]. bioRxiv. 2017. p. 205831. doi:10.1101/205831

26. Friedman J, Higgins LM, Gore J. Community structure follows simple assembly rules in microbial microcosms. Nat Ecol Evol. 2017;1: 109.

27. Gibson TE, Bashan A, Cao H-T, Weiss ST, Liu Y-Y. On the Origins and Control of Community Types in the Human Microbiome. PLoS Comput Biol. 2016;12: e1004688.

28. Swenson W, Wilson DS, Elias R. Artificial ecosystem selection. Proc Natl Acad Sci U S A. 2000;97: 9110–9114.

29. Swenson W, Arendt J, Wilson DS. Artificial selection of microbial ecosystems for 3-chloroaniline biodegradation. Environ Microbiol. 2000;2: 564–571.

30. Williams HTP, Lenton TM. Artificial selection of simulated microbial ecosystems. Proc Natl Acad Sci U S A. 2007;104: 8918–8923.

31. Panke-Buisse K, Poole AC, Goodrich JK, Ley RE, Kao-Kniffin J. Selection on soil microbiomes reveals reproducible impacts on plant function. ISME J. 2015;9: 980–989.

32. Rillig MC, Tsang A, Roy J. Microbial Community Coalescence for Microbiome Engineering. Front Microbiol. 2016;7: 1967.

33. Bairey E, Kelsic ED, Kishony R. High-order species interactions shape ecosystem diversity. Nat Commun. 2016;7: 12285.

34. Levine JM, Bascompte J, Adler PB, Allesina S. Beyond pairwise mechanisms of species coexistence in complex communities. Nature. 2017;546: 56–64.

35. Mayfield MM, Stouffer DB. Higher-order interactions capture unexplained complexity in diverse communities. Nat Ecol Evol. 2017;1: 62.

36. Guo X, Boedicker JQ. The Contribution of High-Order Metabolic Interactions to the Global Activity of a Four-Species Microbial Community. PLoS Comput Biol. 2016;12: e1005079.

37. Sailer ZR, Harms MJ. Detecting High-Order Epistasis in Nonlinear Genotype-Phenotype Maps. Genetics. 2017;205: 1079–1088.

38. Weinreich DM, Delaney NF, Depristo MA, Hartl DL. Darwinian evolution can follow only very few mutational paths to fitter proteins. Science. 2006;312: 111–114.

39. Taylor MB, Ehrenreich IM. Higher-order genetic interactions and their contribution to complex traits. Trends Genet. 2015;31: 34–40.

40. Kuzmin E, VanderSluis B, Wang W, Tan G, Deshpande R, Chen Y, et al. Systematic analysis of complex genetic interactions. Science. 2018;360. doi:H 10.1126/science.aao1729

41. Velenich A, Gore J. The strength of genetic interactions scales weakly with mutational effects. Genome Biol. 2013;14: R76.

42. Sarkisyan KS, Bolotin DA, Meer MV, Usmanova DR, Mishin AS, Sharonov GV, et al. Local fitness landscape of the green fluorescent protein. Nature. 2016;533: 397–401.

43. Poelwijk FJ, Kiviet DJ, Weinreich DM, Tans SJ. Empirical fitness landscapes reveal accessible evolutionary paths. Nature. 2007;445: 383–386.

44. Scholes C, DePace AH, Sánchez Á. Combinatorial Gene Regulation through Kinetic Control of the Transcription Cycle. Cell Systems. Elsevier; 2016;4: 97–108.

45. Buchler NE, Gerland U, Hwa T. On schemes of combinatorial transcription logic. Proc Natl Acad Sci U S A. 2003;100: 5136–5141.

46. Wood K, Nishida S, Sontag ED, Cluzel P. Mechanism-independent method for predicting response to multidrug combinations in bacteria. Proc Natl Acad Sci U S A. 2012;109: 12254–12259.

47. Wood KB, Wood KC, Nishida S, Cluzel P. Uncovering scaling laws to infer multidrug response of resistant microbes and cancer cells. Cell Rep. 2014;6: 1073–1084.

48. Abrams PA. Arguments in Favor of Higher Order Interactions. Am Nat. [University of Chicago Press, American Society of Naturalists]; 1983;121: 887–891.

49. Grilli J, Barabás G, Michalska-Smith MJ, Allesina S. Higher-order interactions stabilize dynamics in competitive network models. Nature. 2017;548: 210–213.

50. Bajić D, Moreno-Fenoll C, Poyatos JF. Rewiring of genetic networks in response to modification of genetic background. Genome Biol Evol. 2014;6: 3267–3280.

51. Summers JW, Wyss O. Biotin-deficient growth of Bacillus polymyxa. J Bacteriol. 1967;94: 1908–1914.

52. Marchesi JR, Adams DH, Fava F, Hermes GDA, Hirschfield GM, Hold G, et al. The gut microbiota and host health: a new clinical frontier. Gut. 2016;65: 330–339.

53. Gomez A, Espinoza JL, Harkins DM, Leong P, Saffery R, Bockmann M, et al. Host Genetic Control of the Oral Microbiome in Health and Disease. Cell Host Microbe. Elsevier; 2017;22: 269–278.e3.

54. Aguilar-Rodríguez J, Payne JL, Wagner A. A thousand empirical adaptive landscapes and their navigability. Nature Ecology & Evolution. Nature Publishing Group; 2017;1: 0045.

55. Inaoka T, Ochi K. Scandium stimulates the production of amylase and bacilysin in Bacillus subtilis. Appl Environ Microbiol. 2011;77: 8181–8183.

56. Fersht A. Structure and Mechanism in Protein Science: A Guide to Enzyme Catalysis and Protein Folding. 1st edition. W. H. Freeman; 1998.

57. Dill KA, Bromberg S. Molecular Driving Forces: Statistical Thermodynamics in Chemistry & Biology. 1 edition. Garland Science; 2002.

58. Abdel-Fattah YR, Soliman NA, El-Toukhy NM, El-Gendi H, Ahmed RS. Production, Purification, and Characterization of Thermostable α-Amylase Produced by Bacillus licheniformis Isolate AI20. J Chem Chem Eng. Hindawi; 2012;2013. doi:10.1155/2013/673173

59. Al-Qodah Z. Determination of kinetic parameters of &#8733-amylase producing thermophile Bacillus sphaericus. Afr J Biotechnol. Academic Journals (Kenya); 2007;6.

